# Segment-specific orientation of the dorsal and ventral roots for precise therapeutic targeting of human spinal cord

**DOI:** 10.1101/2020.01.31.928804

**Authors:** Alan Mendez, Riazul Islam, Timur Latypov, Prathima Basa, Ogeneitsega J. Joseph, Bruce Knudsen, Ahad M. Siddiqui, Priska Summer, Luke J. Staehnke, Peter J. Grahn, Nirusha Lachman, Anthony J. Windebank, Igor A. Lavrov

## Abstract

An understanding of spinal cord functional neuroanatomy is essential for diagnosis and treatment of multiple disorders including, chronic pain, movement disorders, and spinal cord injury. Till now, no information is available on segment-specific spinal roots orientation in humans. In this study we collected neuroanatomical measurements of the dorsal and ventral roots from C2-L5, as well as spinal cord and vertebral bone measurements from adult cadavers. Spatial orientation of dorsal and ventral roots were measured and correlated to the anatomical landmarks of the spinal cord and vertebral column. The results show less variability in rostral root angles compared to the caudal angles across all segments. Dorsal and ventral rootlets were oriented mostly perpendicular to the spinal cord at the cervical level and demonstrate more parallel orientation at the thoracic and lumbar segments. The number of rootlets was the highest in dorsal cervical and lumbar segments. Spinal cord transverse diameter and size of the dorsal columns were largest at cervical and lumbar segments. The strongest correlation was found between the length of intervertebral foramen to rostral rootlet and vertebral bone length. These results could be used to locate spinal roots and spinal cord landmarks based on bone marks on CT or X-rays. These results also provide background for future correlations between anatomy of spinal cord and spinal column structures that could improve stereotactic surgical procedures and electrode positioning for spinal cord neuromodulation.

**One Sentence Summary:** This is the first detailed analysis of the segment-specific dorsal and ventral spinal roots spatial orientation measured and correlated to the anatomical landmarks of the spinal cord and vertebral column for human.

## Introduction

In recent years, spinal cord neuroanatomy has gained great interest, as it directly impact the outcome of epidural electrical stimulation (EES) to improve motor functions after spinal cord injury (SCI) (*1*-*4*) or alleviate chronic pain (*5*). Several computational studies suggested that dorsal roots and dorsal spinal columns could be considered as a main target for EES (*6*, *7*) and that activation of the dorsal roots fibers depends on orientation of electrical field along the target fibers (*8*), as well as in the curvature of the dorsal roots and the angles between the root fibers and the spinal cord axis (*9*-*12*). On a large animal model, EES of the dorsal root entry zone (DREZ) evoked higher responses in both proximal and distal muscles when stimuli were applied within spinal segments (L1, L2 and L3), compared to when stimuli were applied between the segments (L1-L2, L2-L3 and L3-L4) (*13*). Significant anatomical differences in caudal segments L5-L6, such as proximity of dorsal roots and narrowing of caudal root angle, could also explain the difference in amplitude of motor responses compared to stimulation of more rostral segments (*13*). Overall these and other data suggest that EES-induced motor responses are dependent on exact position of the electrode over the DREZ and on orientation of the fibers in stimulated roots. Until now, only a few attempts have been made to describe the spinal cord segment-specific orientation in human and primary for cervical roots (*14*-*17*). Spinal cord anatomical measurements have been described in different studies with particular focus on DREZ length (*17*-*21*), ventral root exit zone (VREZ) length (*21*), intersegmental space (*17*), segment length (*22*), root length (*17*-*21*), distance across columns (*17-19, 22*), root diameter (*23*), and number of rootlets (*17*-*22*). At the same time, no information is currently available on dorsal and ventral roots orientation across all spinal cord segments. The main goal of this study was to comprehensively illustrate the anatomical features of the spinal cord roots and their variations across spinal segments (C2-L5) in human. The information on dorsal and ventral spinal roots orientation and their segment-specific distribution in correlation with spinal cord measurements and specific bone landmarks is critical for future development of effective navigation approaches for implantation of electrodes and for stereotaxic spinal surgeries. This information will also help to further optimize stimulation protocols, i.e. electrodes selection and positioning on the spinal cord and develop novel computational approaches for spinal cord stimulation (SCS).

## Results

### Spinal cord gross anatomy

The major spinal cord gross anatomical landmarks were measured in this study: DREZ length, segment length at dorsal column and bone entry, inferior articular facet to caudal rootlet distance and intervertebral foramen to rostral and to caudal rootlet distance as illustrated on Fig. 1A. Data (mean, SD) from these anatomical measurements are summarized and presented on Fig. 1B-I. The DREZ length and segment length at dorsal column demonstrated a “bell-shape” distribution with the greater lengths at the thoracic segments (Fig. 1B, C, and E). Segment length at bone entry demonstrated the similar pattern from cervical to upper and mid-thoracic segments that continued to increase further at lower thoracic and lumbar segments (Fig. 1D, E). The inferior articular facet, intervertebral foramen to rostral, and to caudal rootlets lengths demonstrated gradual increase from cervical to lumbar segments with slight increase at the high thoracic segments and prominent increase at lumbar segments (Fig. 1F-I).

**Fig. 1.**
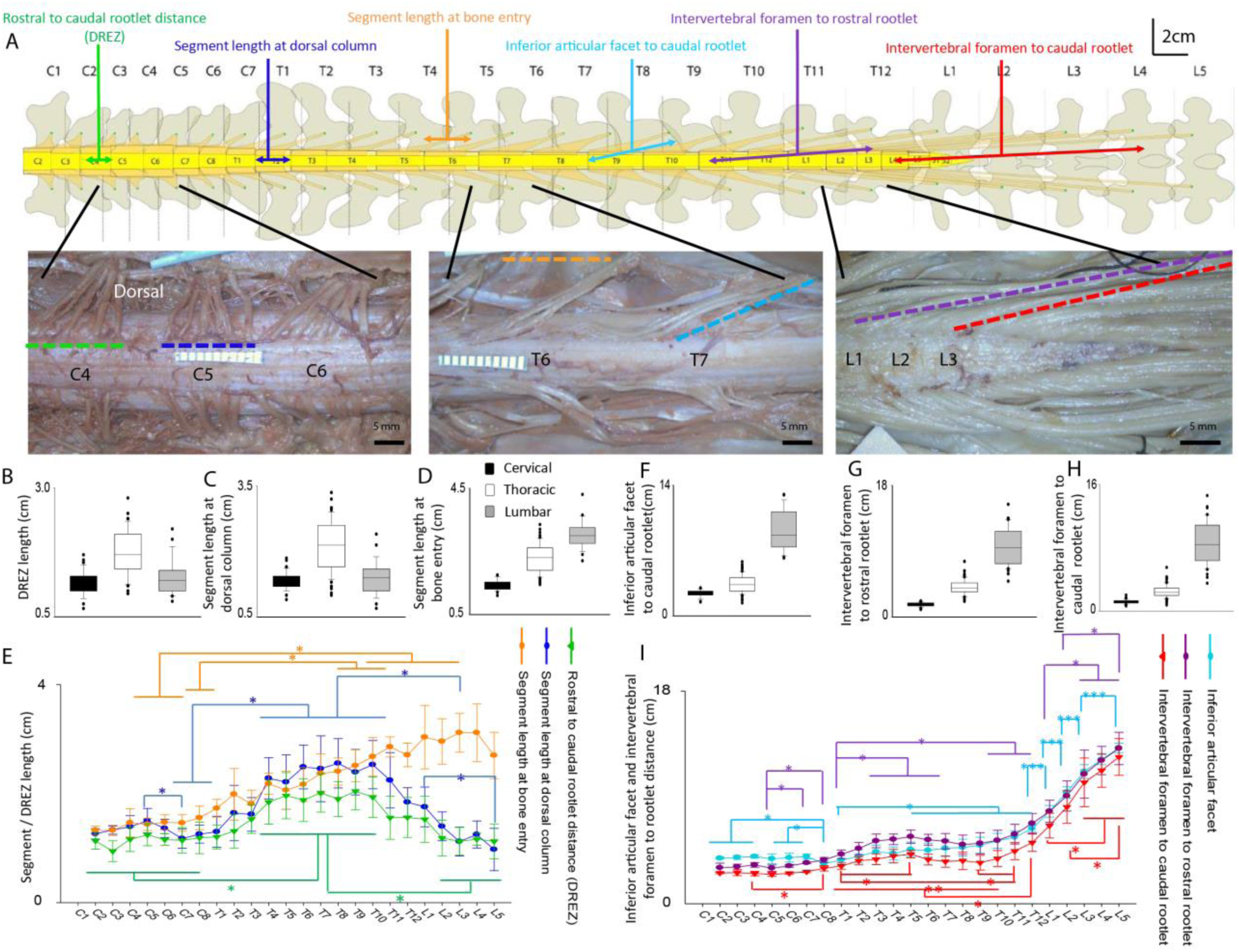
Spinal cord anatomic parameters (**A**). Schematic representation of vertebrae and spinal cord with measured parameters. Examples of dorsal segments correspond to cervical C4, C5, C6; thoracic T6, T7; and lumbar L1, L2 L3, segments. (**B-D**) Bar graphs of DREZ and segments lengths for cervical (black), thoracic (gray), and lumbar (white) segments. (E) Segmental description of DREZ and segments length across C2-L5 segments. Note the similar trend in DREZ length (green line) and segment length at dorsal column entry (blue line), although, segment length was overall higher across all segments except the lower lumbar segments (L3-L5), where no intersegmental space was observed. Segment length at bone entry (orange line) demonstrates consistent increase across all segments. (**F-H**) Bar graphs for inferior articular to caudal rootlet distance and intervertebral foramen to rostral/caudal rootlet distances for cervical (black), thoracic (gray), and lumbar (white) segments. (**I**) Segmental description of inferior articular facet to caudal rootlet distance (light blue line), intervertebral foramen to rostral rootlet distance (purple line), and intervertebral foramen to caudal rootlet distance (red line) for C2-L5 segments show a similar trend across all segments. Data represent mean+/- STD, n=9, One-way ANOVA followed by Kruskal-Wallis, (*p<0.05, **p<0.01 & ***p<0.001)

#### DREZ length

DREZ length was greatest at thoracic segments (1.73 ± 0.42 cm) (p<0.001) (Fig. 1B). Segmental description showed that C2 segment was 1.13 ± 0.15 cm and remained mainly unchanged throughout cervical and upper thoracic segments. DREZ length increased significantly at T4 (1.84 ± 0.40 cm) compared to cervical segments and T1, and did not change significantly through the rest of thoracic segments including L1; however at L2 (1.18 ± 0.32 cm), there was a significant decrease, with no other major variations across the rest of the lumbar segments (Fig. 1E, green line).

#### Segment length at dorsal column entry

Segment length at dorsal column entry was greatest at thoracic segments (2.13 ± 0.57 cm) (p<0.001) (Fig. 1C). Segmental description demonstrated that C2 was 1.26 ± 0.09 cm and did not change significantly until C7 (1.17 ± 0.17 cm), where it decreased. No major variations were observed across low cervical and upper thoracic segments, however, length significantly increased at T4 (2.28 ± 0.37 cm) compared to T1 segment. The segment length remained similar throughout all thoracic segments and upper lumbar segments, although at L3 (1.12 ± 0.26 cm) it decreased significantly compared to T11 and then decreased again at L5 (0.97 ± 0.38 cm) compared to L1 (Fig. 1E, blue line).

#### Segment length at bone entry

Segment length at bone entry was gradually increased over segments and was greatest at lumbar segments (3.01 ± 0.44 cm) (p<0.001) (Fig. 1D). Segmental description showed that C2 was 1.31 ± 0.10 cm and did not change significantly through all cervical segments and upper thoracic segments. A significant increase in segment length was observed at T9 (2.18 ± 0.19 cm), when compared to T1. No major variation was observed in the rest of spinal cord segments (Fig. 1E, green line).

#### Inferior articular facet to caudal rootlet distance

Inferior articular facet to caudal rootlet distance was gradually increased across segments and was greatest at lumbar segments (9.16 ± 2.15 cm) (p<0.001) (Fig. 1F). Segmental description showed that distance at C2 segment was 2.44 ± 0.08 cm and it did not show major variations across all cervical segments, except for C8 (1.94 ± 0.27 cm) where it decreased. This distance did not vary at T1 and T2 segments, but significantly increase at T3 segment (3.03 ± 0.59 cm) compared to C8. No major variations were observed across mid-cervical segments, until at T7 (3.36 ± 0.77 cm), where it showed a significant increase. For the rest of thoracic segments inferior articular facet to caudal rootlet distance remained mainly unchanged. At lumbar segments L1 (6.81 ± 0.55 cm), L2 (8.58 ± 0.51 cm), and L3 (10.52 ± 0.64 cm) it was significantly longer compare to T12. Segments L2 and L3 had longer inferior articular facet to caudal rootlet distance compared to L1. At segments L3 and L4 it did not vary significantly and L5 (12.71 ± 0.44 cm) had greater distance compare to L3 (Fig. 1I, blue line).

#### Intervertebral foramen to rostral rootlet distance

Similar to previous measurement, intervertebral foramen to rostral rootlet distance was gradually increased to the greatest values at lumbar segments (9.59 ± 2.55 cm) (p<0.001) (Fig. 1G). This distance at C2 segment was 1.55 ± 0.42 cm and it did not change significantly throughout cervical segments till C7, where C7 (2.01 ± 0.08 cm) and C8 (2.27 ± 0.22 cm) were significantly greater compare to C5. Lower cervical and upper thoracic segments did not vary significantly till T3, where there it increased (4.05 ± 0.63 cm). No other major variations were observed across the rest of thoracic segments except for T12 (5.68 ± 0.92 cm) that was significantly longer compare to T1-T10. T12, L1, and L2 segments did not vary, but L3 (10.27 ± 2.23 cm) showed significant increase compared to L1. No major variations were observed for the rest of lumbar segments, although at L5 (12.66 ± 1.50 cm) it was significantly longer than at L2 (Fig. 1I, purple line).

#### Intervertebral foramen to caudal rootlet distance

Similar, intervertebral foramen to caudal rootlet distance gradually increased to the greatest values at lumbar segments (8.6 ± 2.76 cm) (p<0.001) (Fig. 1H). This distance of C2 segment was 1.06 ± 0.07 cm and remained relatively unchanged across all cervical segments except for C8 (1.50 ± 0.25 cm), where it was longer compare to other cervical segments. No variations were observed at the upper thoracic segments, although, T5 showed some increase and was significantly longer compare to T1 (2.81 ± 0.35 cm). Intervertebral foramen to caudal rootlet distance increased again at T11, being longer compared to T9 (3.10 ± 0.80 cm). No variations were observed between lower thoracic (T10-T12) and upper lumbar segments (L1-L2). Distance at L3 segment (9.51 ± 2.27 cm) significantly increased compared to L1, and remained similar through the rest of lumbar segments (Fig. 1I, red line).

### Anatomy of the Dorsal and Ventral Root and Rootlets

Main anatomical measurements of the dorsal and ventral roots and rootlets are illustrated in Fig. 2A. Rostral and caudal root angles, number of dorsal and ventral rootlets, width across dorsal and ventral columns, and transverse diameter of dorsal and ventral roots were measured in this study. Data (mean, SD) from these anatomical measurements are summarized and presented on Fig. 2B-P. Most of these measurements had a bimodal (double-peaked) distribution across spinal segments with the higher values at cervical and lumbar segments compared to thoracic segments (i.e. number of roots, root diameter, and transversal width). A reverse bimodal distribution was found for dorsal and ventral width (Fig. 2M). A continuous increase in values across all segments was observed for rostral root angle, while continuous decrease was consistent for caudal root angles (Fig. 2F).

**Fig. 2.**
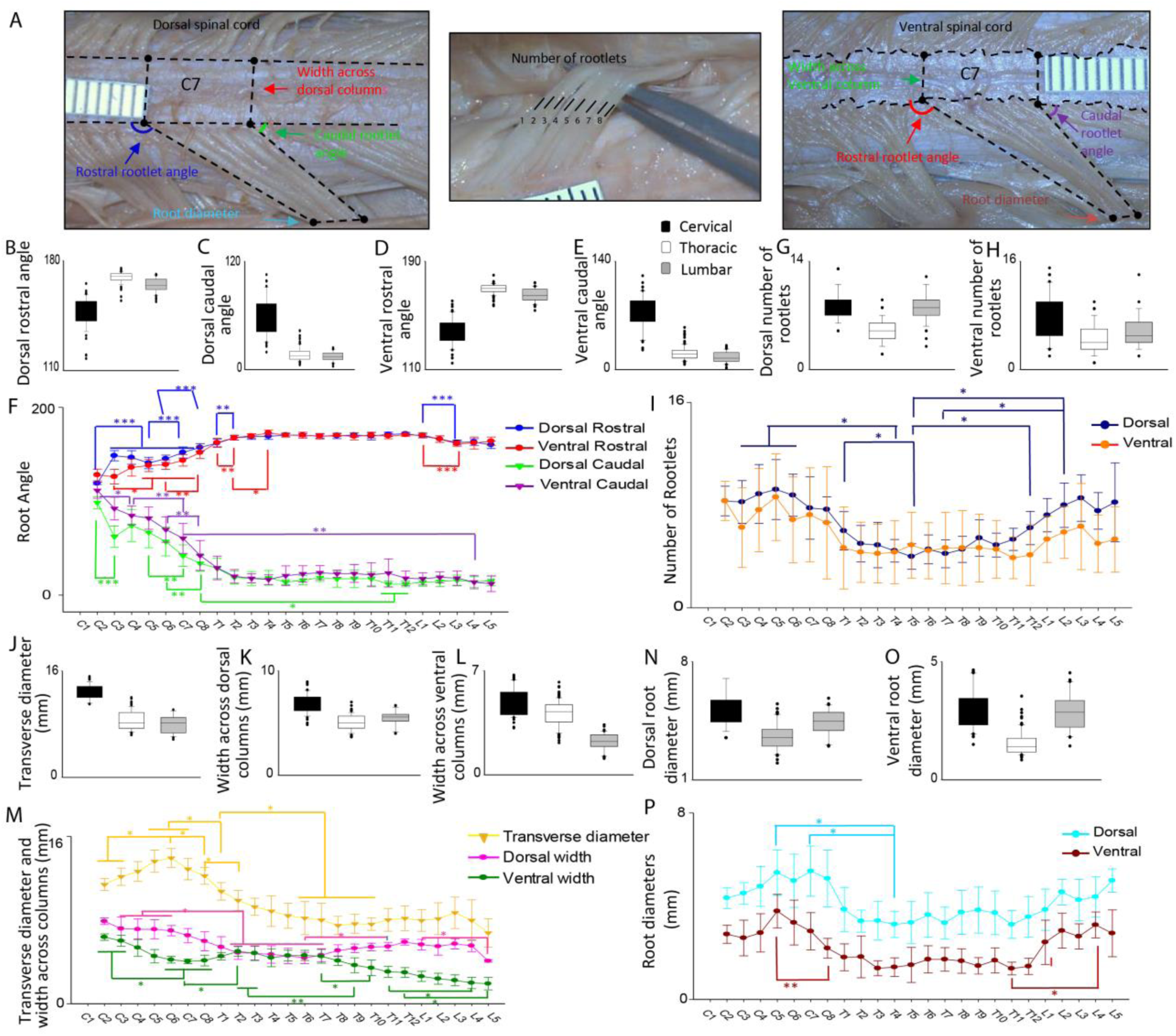
Roots and rootlets anatomic parameters (**A**) Dorsal and ventral roots and rootlets anatomic measurements. Scales in images represent millimeters. (**B-E**) Bar graphs for dorsal and ventral roots orientation (angles) for cervical (black), thoracic (gray), and lumbar (white) segments. (**F**) Segmental description of dorsal and ventral roots angles for C2-L5 segments. Dorsal and ventral roots angles follow the same orientation when entering the spinal cord. Note the trend of increase from dorsal rostral (blue line) and ventral rostral (red line) angles; opposed to the decrease in dorsal caudal (green line) and ventral caudal (purple line) angles. (**G-H**) Bar graphs for dorsal and ventral number of rootlets for cervical (black), thoracic (gray), and lumbar (white) segments. (**I**) Segmental description of number of dorsal (dark blue line) and ventral (orange line) rootlets. (**J-L**) Bar graphs for spinal cord transverse diameter, and dorsal and ventral widths across columns for cervical (black), thoracic (gray) and lumbar (white) segments. (**M**) Segmental description of spinal cord transverse diameter (yellow line), and dorsal (pink line) and ventral (dark green line) widths across columns for C2-L5 segments. (**N-O**) Bar graphs for dorsal and ventral roots diameter for cervical (black), thoracic (gray) and lumbar (white) segments. (**P**) Segmental description of dorsal (cyan line) and ventral (brown line) roots diameter for C2-L5 segments. (Data represent mean+/- STD, n=9, One-way ANOVA followed by Kruskal-Wallis, *p<0.05, **p<0.01 & ***p<0.001)

#### Rostral roots angles

Dorsal rostral root angle was the greatest at thoracic segments (169.21 ± 3.58°) (p<0.001) (Fig. 2B). At C2 dorsal rostral root angle was 119.05 ± 1.27° and increased significantly at C3 to 148.51 ± 5.43° and then remained consistent throughout the mid-cervical segments. Dorsal rostral root angle increased again at C7 (151.99 ± 5.96°) compared to C5 (140.78 ± 6.12°) and at the C8 segment (157.31 ± 4.06°) where it was significantly greater compare to C6 (145.20 ± 3.86°). At the thoracic segments, there was an increase from T1 to T2 (T1: 161.96 ± 4.42°, T2: 167.60 ± 2.44°), but then it remained unchanged throughout rest of the thoracic and upper lumbar segments. Rostral root angles then decreased significantly at L3 (162.68 ± 2.61°) compared to L1 (169.54 ± 2.04°) (Fig. 2F). For ventral side, rostral angle was also greatest at thoracic segments (169.42 ± 3.63°) (p<0.01) (Fig. 2D). At C2 ventral rostral root angle was 128.03 ± 6.23° and remained constant through C3-C4. It significantly increased at C5 (138.08 ± 6.54°) compared to C3 and did not change significantly at the mid-cervical level, although, it increased at C8 (151.86 ± 6.42°) compared to C6. Significant increase in ventral rostral root angle was observed at T2 (167.76 ± 2.70°) and then again at T4 (172.82 ± 3.03°). Then, it remained consistent across the rest of thoracic segments. At the lumbar segments the ventral rostral root angle decreased at L3 (161.27 ± 2.74°) and remained similar through the rest of lumbar segments (Fig. 2F).

#### Caudal roots angles

Dorsal caudal root angle was greatest at cervical segments (58.05 ± 21.29°) (p<0.001) (Fig. 2C). At C2 dorsal caudal root angle was 98.57 ± 6.35°. It markedly decreased at C3 (62.17 ± 11.21°) with no major variations across the mid-cervical segments. Then, it decreased again at low cervical level where at C7 (41.94 ± 11.47°) it was smaller compare to C5, and then decreased again at C8 (33.76 ± 8.36°) compared to C6. Then, it remained unchanged throughout most of thoracic segments and decreased again at T11 (11.67 ± 3.06°). No major variations were observed throughout T12 and the lumbar segments (Fig. 2F). For ventral side, ventral caudal root angle was also greatest at cervical segments (73.901 ± 22.7°) (p<0.001) (Fig. 2E). At C2 ventral caudal root angle was similar to C3 (111.55 ± 8.37°), however, it significantly decreased at C4 (84.81 ± 10.00°) and then again decreased at C7 (60.48 ± 16.03°). Caudal root angles remained similar throughout thoracic and lumbar segments till L4 (13.74 ± 7.11°) where they decreased significantly compared to C7 (Fig. 2F).

#### Number of Rootlets

As for dorsal roots, cervical and lumbar segments demonstrated the greatest number of rootlets (8.41 ± 1.91 and 7.91 ± 1.94 respectively) (p<0.001) (Fig. 2G). The number of rootlets at cervical C2 segment was 8.33 ± 1.15 and remained relatively similar throughout all cervical and upper thoracic segments. It significantly decreased at T4 (4.44 ± 0.88) compared to mid-cervical segments (C4-C6) and remained unchanged throughout the rest of the thoracic segments. There was a significant increase in rootlet number at L2 (8.00 ± 1.73) that was greater compare to T5 and T7, without major variations in the rest of lumbar segments (Fig. 2I). Ventral roots demonstrated bimodal distribution with the greatest average number of rootlets at cervical and lumbar segments (7.28 ± 3.35 and 5.57 ± 2.38 respectively) (p<0.05) (Fig. 2H). Although there was a trend for increased number of rootlets in cervical and lumbar segments, no statistical significance was found (Fig. 2I). Thus, the increase in number of rootlets in cervical and lumbar segments was only evident for dorsal roots that could be related to the functional role of the sensory information for forelimb and hind limbs represented across these segments (Fig. 2I).

#### Transverse diameter

Since no statistical differences were found between the three measured transverse diameters at DREZ with respect to rostral, middle or caudal reference points (data not shown), we averaged them into a single diameter measurement per each segment. Transverse diameter was the greatest at cervical segments (12.86 ± 1.13 mm) (p<0.001) (Fig. 2J). Segmental description showed that C2 transverse diameter was 11.4 ± 0.5 mm and remained significantly unchanged until C6, where it demonstrated significant increase (13.9 ± 0.8 mm) and then decreased at C8 (12.2 ± 0.7 mm) compared to C6. At C8 and T1 transverse diameters were not significantly different, although, transverse diameter decreased significantly at T2 (9.8 ± 0.9 mm) and then decreased again at T6 (8.1 ± 1.5 mm). Then, transverse diameter remained consistent throughout rest of the thoracic and lumbar segments (Fig. 2M). Although there was a trend with larger diameter at the lumbar segments, the only statistically significant increase was found at the cervical segments.

#### Width across dorsal and ventral columns

Width across dorsal columns was greatest at cervical segments (6.84 ± 0.93 mm) (p<0.001) and the smallest at the thoracic segments (5.05 ± 0.72 mm) (p<0.05) (Fig. 2K). At C2 dorsal columns width was 7.88 ± 0.30 mm and it did not change significantly across all cervical segments and T1. Then, at T2 (4.87 ± 0.63 mm) dorsal columns width decreased and remained largely unchanged throughout mid-thoracic segments, and then increased at T11 (5.45 ± 0.39 mm) compared to T6. The dorsal columns width remained unchanged throughout T12 and most of lumbar segments, except for L5 (4.08 ± 0.11 mm), where it demonstrated significant decrease compared to L1 and L3 (Fig. 2M). Width across ventral columns similar had greatest values at cervical segments (4.78 ± 0.94 mm) (p<0.05), although, opposite to dorsal columns width, the smallest ventral columns width was found at the lumbar segments (2.22 ± 0.56 mm) (p<0.001) (Fig. 2L). At C2 width across ventral columns was 6.36 ± 0.30 mm and did not change till C6 (4.20 ± 0.31 mm) where it decreased significantly compared to C2 and C3. No major variations were observed at lower cervical segments and T1 till T2 (5.00 ± 0.46 mm) where it was greater compared to C7. Width across ventral columns remained consistent through mid-thoracic segments and, then decreased again at T9 (3.67 ± 0.71 mm) segment, where it was significantly smaller compared to T2 and T3. At T10 (3.42 ± 0.74 mm) it was smaller compare to T1-T7. At T12 (2.98 ± 0.46 mm) ventral columns width was smaller compare T1-T8. Ventral columns width did not vary significantly at the upper lumbar segments, although, it demonstrated significant decrease at L4 (1.99 ± 0.54 mm) being smaller compare to T11 (Fig. 2M). Inverted bimodal distribution was observed in dorsal vs. ventral width, indicating that at cervical and lumbar segments have relatively wider dorsal columns compare to shorter columns width at ventral side (Fig. 2M).

#### Root diameter

For the dorsal side, the greatest root diameters were observed at cervical and lumbar segments (5.05 ± 0.98 mm and 4.38 ± 0.8 mm respectively) (p<0.001) (Fig. 2N). At C2 dorsal root diameter was 4.34 ± 0.45 mm and significant variations were only observed at segments C5 (5.43 ± 0.98 mm) and C7 (5.50 ± 1.08 mm), both being significantly larger than T4 (Fig. 2P). For the ventral side, cervical and lumbar segments also demonstrated the greatest diameters (2.94 ± 0.86 mm and 2.81 ± 0.78 respectively) (p<0.001) (Fig. 2O). At C2 ventral root diameter was 2.79 ± 0.38 mm and remained unchanged till C8 (2.19 ± 0.42 mm), where it decreased. Then, ventral root diameter remained the same across all thoracic segments and upper lumbar segments. A marked increase in ventral root diameter was observed at L4 (3.19 ± 0.55 mm), without major change at L5 (Fig. 2P).

### Related spine anatomical landmarks

The bone anatomical landmarks were measured as illustrated on Fig. 3A: mid-vertebrae foramen length, vertebrae bone length, intervertebral foramen distance, intervertebral foramen diameter. The data (mean, SD) from these anatomical measurements are summarized and presented on Fig. 3B. Most of these spinal anatomical measurements demonstrated gradual increase from cervical to lumbar segments with slight increase at mid-cervical and mid-lumbar segments.

**Fig. 3.**
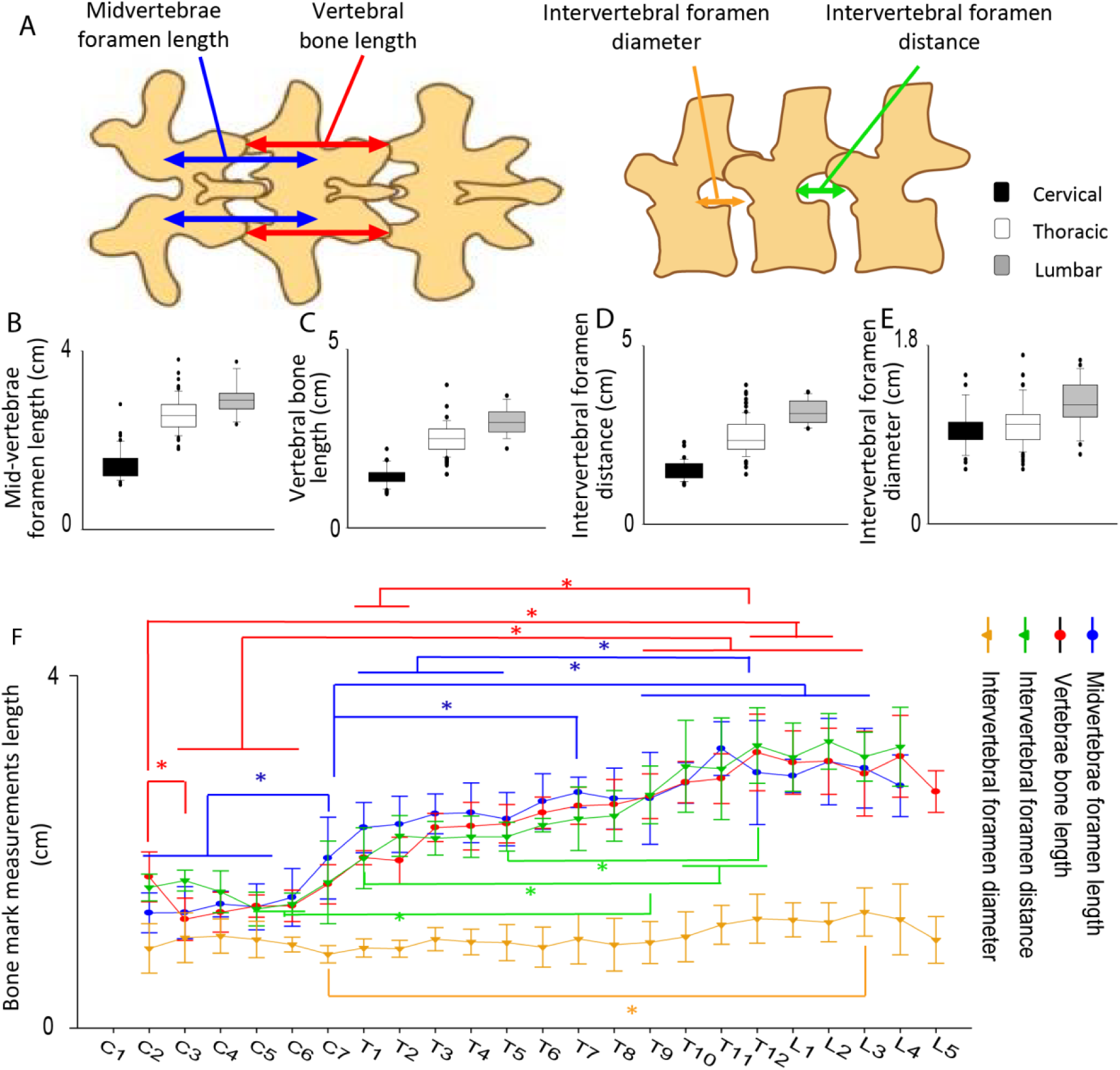
Vertebral anatomic landmarks (**A**) Spine anatomic landmarks (postero-anterior and lateral view). (**B-E**) Bar graphs for mid-vertebrae foramen length, vertebral bone length, intervertebral foramen distance and intervertebral foramen diameter for cervical (black), thoracic (gray), and lumbar (white) segments. (**F**) Mid-vertebrae foramen length (blue line), vertebral bone length (red line), intervertebral foramen distance (green line) and intervertebral foramen diameter (yellow line) for C2-L5 segments. Note the trend equivalence in mid-vertebral foramen length, vertebrae bone length and intervertebral foramen distances as intervertebral measurements and vertebral size increase towards lower levels in a gradual pattern. (Data represent mean+/- STD, n=9, One-way ANOVA followed by Kruskalwalis, *p<0.05, **p<0.01 & ***p<0.001)

#### Mid-vertebrae foramen length

The greatest mid-vertebrae foramen lengths were observed at thoracic and lumbar segments (2.55 ± 0.39 cm and 2.9 ± 0.36 cm respectively) (p<0.001) (Fig. 3B). Segmental description showed that length at C2 vertebra was 1.30 ± 0.22 cm and did not vary significantly between cervical vertebras except for C7 (1.92 ± 0.46 cm), which showed a significant increase. No major variations were observed between lower cervical, upper and mid-thoracic vertebra, however, T7 (2.67 ± 0.17 cm) showed a significant length increase compared to C7. No major variations were observed between rest of vertebrae, although T11 (3.17 ± 0.30 cm) showed to be significantly longer than T1-T5 (Fig. 3F).

#### Vertebral bone length

Vertebrae bone length was greatest at lumbar segments (2.96 ± 0.38 cm) (p<0.001) (Fig. 3C). In segmental description we observed that C2 vertebral length was 1.71 ± 0.28 cm, decreasing at the C3 (1.23 ± 0.24 cm) vertebrae, then, length remained mainly unchanged throughout the rest of cervical and upper thoracic vertebrae. At T9 (2.63 ± 0.25 cm), however, there was a significant increase compared to C6. There was another significant increase at T12 (3.12 ± 0.43 cm) vertebrae, which was significantly longer than T1 and T2 (Fig. 3F). No other major variations were observed at lumbar vertebras.

#### Intervertebral foramen distance

Intervertebral foramen distance showed to be greatest at lumbar segments (3.14 ± 0.33 cm) (p<0.001) (Fig. 3D). Segmental description showed that distance for C2 segment was 1.59 ± 0.15 cm and remained consistent throughout rest of the cervical segments and the upper- and mid-thoracic vertebrae until T9 (2.64 ± 0.32 cm) where a significant increase was observed compared to C5 and C6. There was another significant increase at T10 (2.97 ± 0.52 cm) and T11 (2.94 ± 0.57 cm) compared to T1, and again increased at T12 (3.20 ± 0.42 cm), which was significantly greater compared to T5 (2.17 ± 0.16 cm) (Fig. 3F). No other significant variations were observed in the remainder of the vertebrae.

#### Intervertebral foramen diameter

Intervertebral foramen diameter showed to be greatest at lumbar segments (1.21 ± 0.25 cm) (p<0.001) (Fig. 3E). Segmental description showed that diameter for C2 was 0.90 ± 0.28 cm and remained mainly unchanged across all vertebrae. The only difference observed was at L3 (1.31 ± 0.27 cm) where the diameter was significantly greater than C7 (0.83 ± 0.09 cm) (Fig. 3F).

### Other findings

#### DREZ vs. VREZ

In this study we found distinct differences between dorsal and ventral aspects of the spinal cord. Particular differences across all the spinal segments were found in the way the root fibers enter or exit the spinal cord. Dorsal root fibers at DREZ entered dorsal columns in a well-organized linear fashion, allowing visualization of the posterolateral sulcus as a single line throughout all spinal segments (Fig. 4, Dorsal). For ventral roots at VREZ, we observed root fibers exiting the ventral columns in a less organized fashion, without a clear demarcation line (Fig. 4, Ventral). Another difference is that ventral fibers produced multiple branches compare to dorsal rootlets (Fig. 2 and 4).

**Fig. 4.**
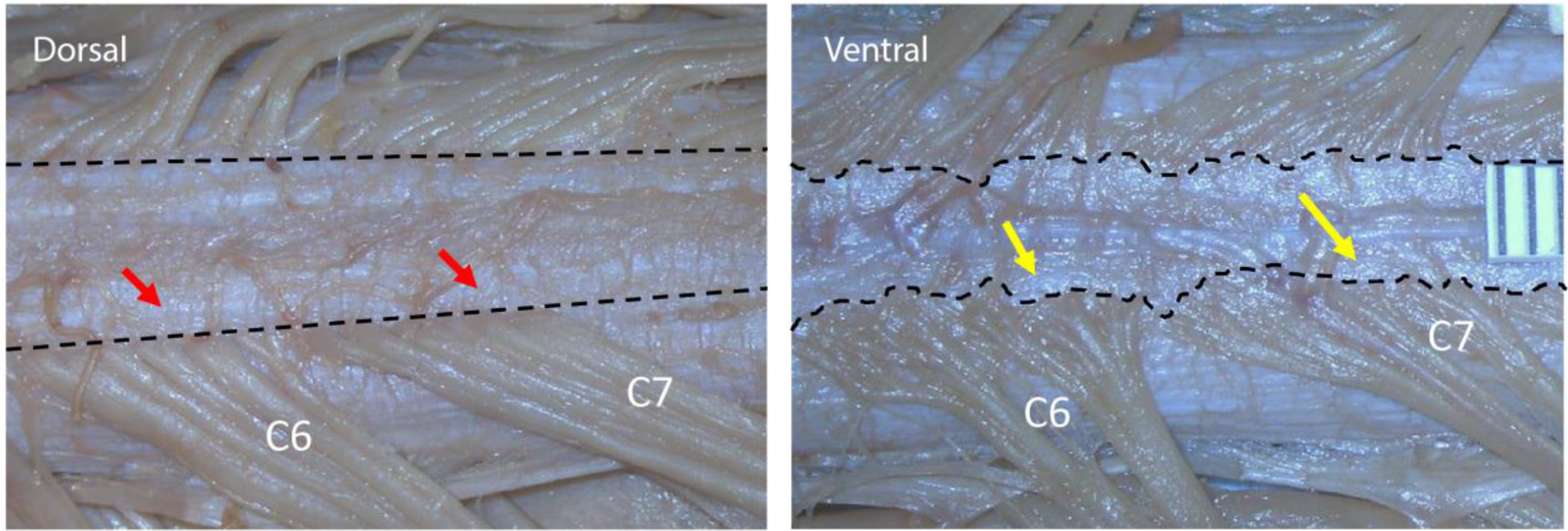
Dorsal and ventral root emergence. Dorsal roots nerve fibers enter dorsal columns in a well demarcated, organized, singular linear fashion (red arrow), following the same pattern through the all spinal cord segments. On the contrary, Ventral root nerve fibers exit ventral columns in a less organized, irregular fashion with multiple branching (yellow arrow). Both images correspond to the dorsal and ventral aspects of C6 and C7 segments, a difference consistent for all levels from cervical to lumbar spinal segments.

#### Inter-root anastomoses

Another finding of this study was the presence of intradural dorsal inter-root anastomoses, which were found at cervical and lumbar segments, but not at thoracic segments. A total of 85 inter-root anastomoses were found across 9 cadavers (Fig. 5A). Cervical roots had higher number of anastomoses (77) compared to lumbar roots (8) (Fig. 5B). Cervical anastomoses were more common on the left side (44) compared to the right side (33), same as for lumbar segments, with more anastomoses on the left side (6) compared to the right (2). The most consistent inter-root anastomoses were found on the left side in segments C3-C4, which was confirmed in all 9 cadavers (Fig. 5B). No inter-root anastomoses were found on the ventral side of the spinal cord.

**Fig. 5.**
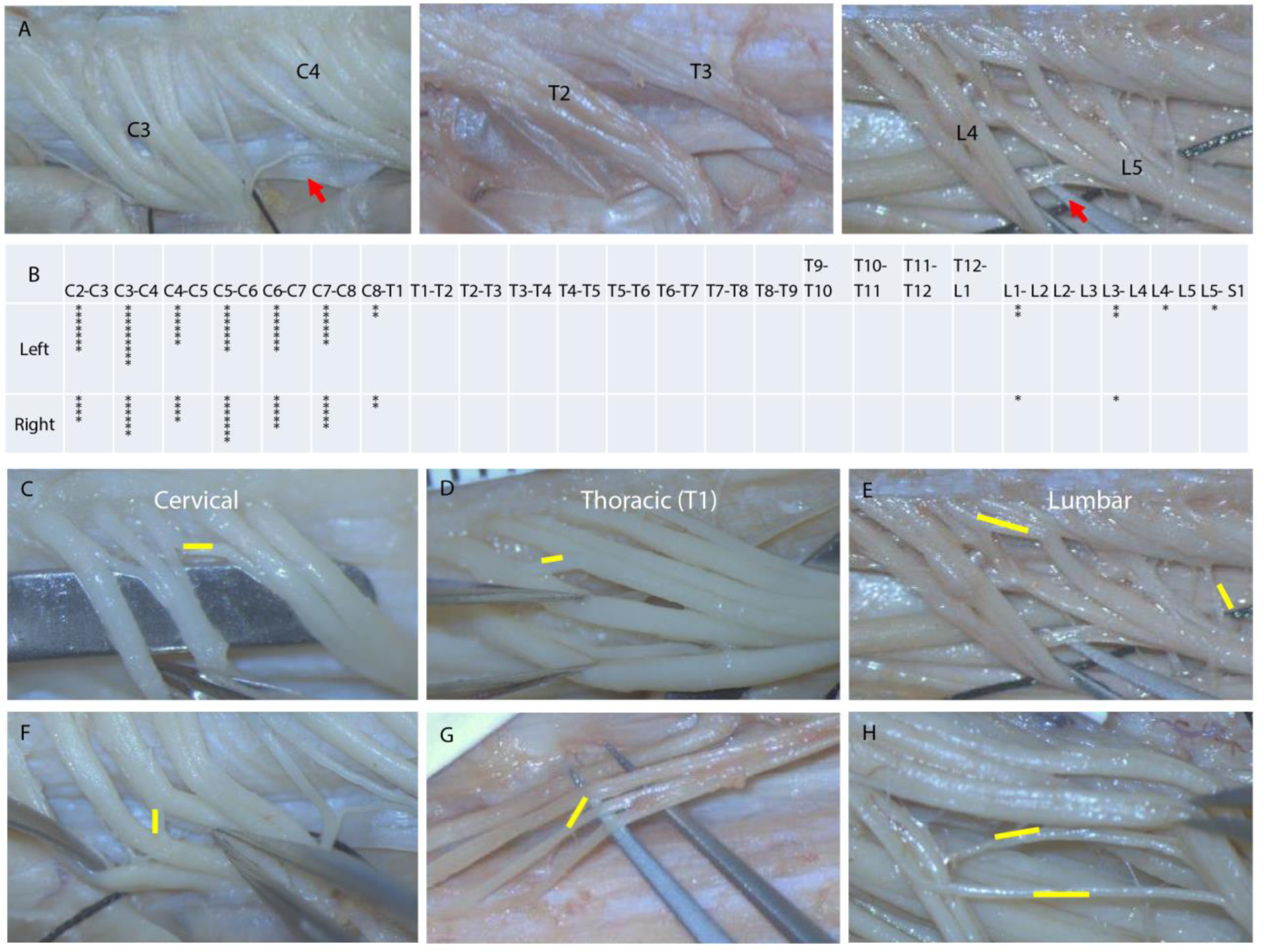
Dorsal inter-root and inter-rootlet anastomoses (**A**) Dorsal inter-root anastomoses (red arrows) corresponds to left dorsal C3-C4, T2-T3, L4-L5 roots. (**B**) Table shows the frequency of left and right inter-root anastomoses for 9 cadavers. Dorsal inter-rootlet anastomoses marked by yellow lines. (**C-H**) Examples correspond to segments at cervical (C3, C5), thoracic (T1), and lumbar (L4, L5) levels.

#### Inter-rootlet anastomoses

Inter-rootlet anastomoses were also observed, and consisted of nerve fibers connecting dorsal rootlets inside of the root. They were found primary at cervical and lumbar segments, and at thoracic T1 segment, although, no inter-rootlet anastomoses were observed at the rest of the thoracic segments (T2-T12) (Fig. 5C-H). Size, orientation, and number of inter-rootlet anastomoses varied from segment to segment and from subject to subject.

#### Correlation between anatomical landmarks of the spine and spinal cord

Correlation analysis between spine and spinal cord was performed in order to establish an anatomical segmental relationship and optimize targeting the spinal structures based on vertebral bone landmarks. Spearman correlation analysis was used to establish relationship between the following spinal cord and spine anatomical landmarks: intervertebral foramen to rostral/caudal rootlet distance, mid-vertebral foramen length, vertebral bone length, and intervertebral foramen distance for all measured spinal cord segments (C2-L5). A stronger relationship was found between vertebral measurements and intervertebral foramen to rostral rootlet distance compared to intervertebral foramen to caudal rootlet distance. The strongest correlation was found between intervertebral foramen to rostral rootlet distance and vertebral bone length (R = 0.82) (p<0.001) (Fig. 6D), whereas intervertebral foramen to rostral rootlet distance and intervertebral foramen distance was weaker (R = 0.78) (p<0.001), and intervertebral foramen to rostral rootlet distance to mid-vertebral foramen length had the weakest correlation (R = 0.73) (p<0.001). For caudal rootlet distance, intervertebral foramen to caudal rootlet distance and intervertebral foramen distance had the strongest correlation (R = 0.76) (p < 0.001), whereas intervertebral foramen to caudal rootlet distance to vertebral bone length (R = 0.75) (p < 0.001) (Fig. 6E) and to mid-vertebral foramen length had weaker correlations (R = 0.70) (p < 0.001). A schematic representation of the spine and spinal cord for all measured segments (C2-L5) was performed by plotting vertebral bone lengths starting from L5 and presented on Fig. 1A. Intervertebral foramen diameters were used to space vertebras. Rostral and caudal root angles were used to determine root orientation. To determine vertical lengths of segments, DREZ lengths and segments lengths at dorsal column entry were plotted (Fig. 6C). Analyzed correlations were used to corroborate the accuracy in the location of each segment by determining the relationship between each vertebra to its corresponding spinal segment. The strongest correlation (intervertebral foramen distance to rostral rootlet distance to vertebral bone length) was used to establish segmental relationship between vertebral bone landmarks and spinal cord (Fig. 6C, D, E).

**Fig. 6.**
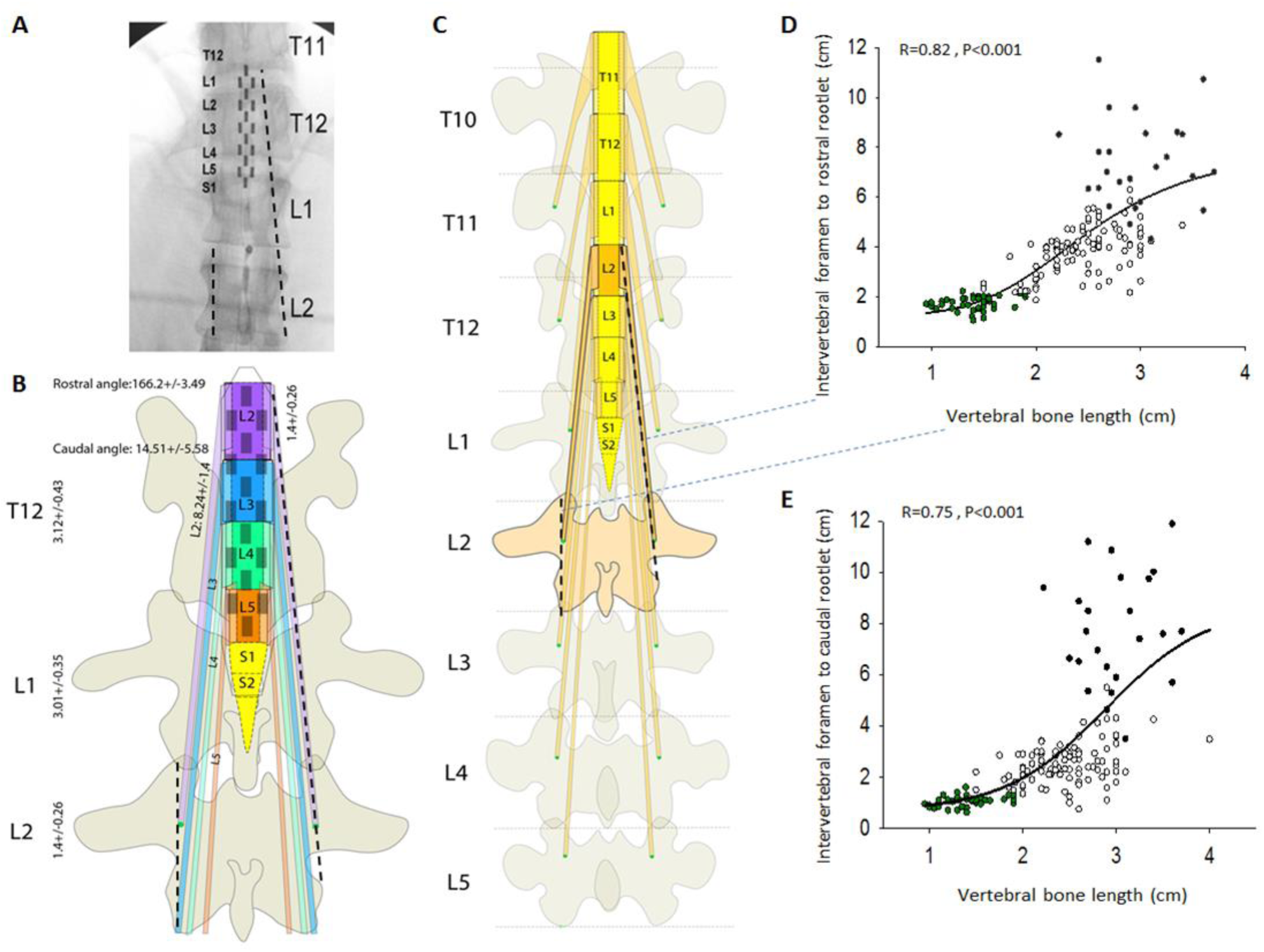
Vertebral spine and spinal cord anatomic relationship for segment mapping and spinal array navigation. (**A**) Fluoroscopy image of T11, T12, L1 and L2 vertebras with related spinal segments (*image adapted from Calvert el al. (24*)). (**B**) Diagram of vertebras (T12-L2) and corresponding spinal cord segments (L2-S2) with array location based on segment mapping. (**C**) Diagram of vertebras (T10-L5) and corresponding spinal cord segments (T11-S2); dotted lines mark L2 intervertebral foramen to rostral rootlet distance and vertebral bone size, parameters that showed strongest correlation. (**D**) Scatter plot of vertebral bone length vs intervertebral foramen to rostral rootlet distance. Spearman coefficient of correlation value was found to be 0.82 with P<0.001. The equation used to fit the curve was, f= 8.5/(1+exp(-(x-2.4)/0.75)). (**E**) Scatter plot of vertebral bone length vs intervertebral foramen to caudal rootlet distance. Spearman coefficient of correlation value was found to be 0.75 with P<0.001. The equation used to fit the curve was, f= 7.5/(1+exp(-(x-2.88)/0.54)).

## Discussion

Multiple computational (*9*-*11*) and animal studies (*13*, *25*) demonstrated that dorsal roots and dorsal columns are the main target for EES, which emphasizes the importance of electrodes location in relation to the main target structures (*9*-*12*). It has been reported that electrical stimulation applied at DREZ provides significantly higher motor evoked responses compared to other electrodes location on dura mater (*3*, *13*). Orientation of electrical field along the fibers of target structures is another key factor in effect of EES (*8*). Surprisingly, up to now no information is available on variation in segment-specific anatomy of dorsal and ventral roots for humans. The segment-specific orientation of spinal roots as well as main anatomical features of the human spinal cord remains virtually unknown, except a few studies that reported results on cervical roots orientation (*14*-*17*). To our knowledge this work is the first attempt to provide comprehensive anatomical measurements of the spinal cord from cervical to lumbar segments, dorsal and ventral roots, their dimensions, and orientation in relation to other spinal cord structures and to vertebral bone marks in humans.

### Dorsal and ventral roots and rootlets neuroanatomy

Anatomical measurements of the human spinal cord have been partially presented in previous works (*14*-*22*). Similar to these studies, we found that length of the dorsal roots, measured as a distance between intervertebral foramen to rostral/caudal rootlet, at cervical segments was shorter compare to thoracic and lumbar (p<0.001) (Fig. 1I). Also, dorsal roots at cervical segments had more perpendicular orientation compared to thoracic and lumbar segments (p<0.001) (Fig. 2F). The largest spinal cord transverse diameter was found at cervical segments (p<0.05) with the maximum at C6 (1.39 ± 0.08 cm) (p<0.05). Similarly, dorsal columns width was the greatest at cervical segments (p<0.001) and smallest at thoracic segments (p<0.05). In opposite, ventral column width was the greatest at the thoracic segments and the smallest at the lumbar segments (p<0.001). Previous studies have analyzed dorsal widths separately for cervical (*17*, *18*), thoracic (*19*), and lumbar segments (*22*) and ventral column width was described only for lumbar segments (*22*) and to our knowledge, this is the first attempt to measure and compare the dorsal and ventral columns widths across cervical, thoracic, and lumbar segments on the same subjects.

### Variation in spinal cord gross anatomy

In this study we observed a bimodal distribution of transverse spinal cord diameter and dorsal column width and inverse bimodal distribution of ventral column width. Thus, dorsal columns at cervical and lumbar segments have a higher width that was the opposite for ventral column width (Fig. 2M). This specific distribution could be related with a number of the dorsal root fibers entering the spinal cord at cervical and lumbar segments responsible for sensory information coming from the lower and upper extremities. This distribution could be also related with the number of roots fibers and was correlated with the highest number of rootlets and root diameter at cervical and lumbar segments (Fig. 2I, P). We also found that DREZ was always presented as a clear demarcation line, while VREZ forms an irregular area (Fig. 1A, 4) related with variations in ventral roots branches that could contribute to the narrowing of the ventral column width, which was also mentioned in previous studies (*26*). The most extensive spinal cord linear growth up to adulthood was previously found at the thoracic segments (*27*) and could explain observed in our study bell-type distribution of segments/DREZ length with the longest DREZ length at the thoracic segments (Fig. 1E) (p<0.001). The extensive growth at the thoracic segments could be also related with a minimal number of anastomoses between roots and rootlets at the thoracic segments (*27*). In this study, at the thoracic segments we did not observe any inter-root anastomoses and only a few inter-rootlet anastomoses (Fig. 5). Formation of inter-root anastomoses, which represent myelinated fibers connecting dorsal roots, has been related to undivided neural crest tissue along the spinal cord (*27*-*29*). This anatomical connectivity may explain overlapping of the sensory symptoms caused by nerve root compression (*29*). As previously described, dorsal roots generally form slower compare to the ventral roots and do not start to separate until approximately the 30th day of gestation (*28*). This could explain the absence of anastomoses between ventral roots, as observed in this and in other studies (*28*-*31*). Our results also show that the greatest number of dorsal inter-root anastomoses at cervical segments (90.58%) compared to other segments (Fig. 5), which may indicate the functional role of these anastomoses in processing the information to the upper extremities (*28*, *32*). Interestingly, we found the greater number of inter-root anastomoses on the left (58.8%) compared to the right side, which has been hypothesized to be related to Lefty-1 gene expression on the left of the neural tube (*29*). Inter-rootlet anastomoses have been identified in the past as connections between rootlets located posterior to dorsal ganglions and connect nerve bundles before they enter the spinal cord (*21*). In this study, we found inter-rootlet anastomoses only within dorsal cervical and lumbar segments varied in shape, size, and number (Fig. 5).

### Dorsal and ventral roots orientation

Only a few anatomy studies describe the segment-specific orientation of the dorsal roots and mainly for cervical segments (*14*-*17*) (Table 1).

**Table 1.** Dorsal and ventral rostral root angles comparison

| Author | Cadaver Count | Spinal Segments Analyzed | Angle Measurement | Statistical Difference |
| --- | --- | --- | --- | --- |
| **Rostral root angle for dorsal segments** |  |  |  |  |
| Alleyne *et al.*, 1998 (15) | 10 | C3-T1 | Smallest: C5 (116°)<br>Greatest: T1 (156°) | P < 0.01<br>P < 0.01 |
| Kumar *et al.*, 2014 (17) | 25 | C1-T1 | Smallest: C1 (110.2 ± 8.2°)<br>Greatest: C5 (144.2 ± 5.3°) | -<br>- |
| Present Study | 9 | C2-L5 | Smallest: C2 (119.0 ± 1.2°)<br>Greatest: T12 (171.8 ± 1.5°) | p < 0.001<br>- |
| **Rostral root angle for ventral segments** |  |  |  |  |
| Shinomiya *et al.*, 1994 (14) | 36 | C5-C8 | *Greatest: C5 (30.9 ± 7.6°) | p < 0.05 |
| Present Study | 9 | C2-L5 | Greatest: T4 (172.8 ± 3.0°) | P < 0.05 |

\*Measured at inferior margin of most superior rootlet

In this study, variation in root angles across spinal segments was consistent with progressive widening of the rostral angles for both dorsal and ventral roots throughout spinal segments, while caudal angles demonstrated progressive narrowing (Fig. 2). Cuellar *et al.*, 2017, demonstrated on a swine model a significant narrowing (< 80°) of caudal angles from proximal lumbar (L1-L3) to the distal lumbar segments (L4-L6), and variation of the proximity of dorsal roots in the same fashion (*13*). As these were the main variation in dorsal roots, they were considered to be related to variation in motor responses amplitude observed with SCS at the most caudal compared to the rostral segments. These variations to some extent resemble the morphologic aspects of lower thoracic and lumbar roots in humans as observed in this study: wide rostral root angles (> 150°), narrow caudal root angles (< 30°), and proximity of dorsal roots (shortening of segments).

### Correlation between neuroanatomy of the spinal roots, spinal cord, and vertebrae and clinical applications

The importance of the location of epidural electrodes during SCS has been described in several studies where EES was beneficial in reducing the muscle tone (*33*) or in activation of the spinal cord locomotor networks in humans (*34*). Most of the currently available surgical procedures on the spinal cord involve implantation of the electrodes based on bony landmarks visualized with fluoroscopy, CT, or ultrasound, in order to locate a point of the needle/electrode insertion. This approach, however, could not provide precise location of the electrode in relation to the spinal cord structures, which are the main targets of neuromodulation. Several attempts to correlate spinal cord and vertebrae structures have been made in the past by correlating spinal segments with vertebral bodies, although they did not lead to practical application (*13*, *20*, *35*). In this study, we correlated spinal structures and bone landmarks recognizable on fluoroscopy to improve electrode placement during surgical implantations and surgical procedures related to the dorsal or ventral spinal roots. Correlation conducted based on collected in this study data showed the strongest relationship between intervertebral foramen to rostral rootlet distance and vertebral bone length (R = 0.82, p < 0.001). By using multiple measurements collected in this study, we have reconstructed C2-L5 spinal segments and roots with corresponding vertebras and their variations presented on Fig. 1A, and specifically for lower thoracic and lumbar segments on Fig. 6, the common target for SCS after SCI. Previously, Lang *et al.*, 1983, used segments length for spinal cord mapping and found that L1 and partial L2 segments at T11 vertebral level and L3-L5 segments at T12 vertebral level can be used for correlation between vertebrae and spinal cord structures (*35*). In another study, authors marked intervertebral disks between T10 to L2 vertebras and performed cross-sections of the roots above each intervertebral disk to map corresponding segments and found that L1-L3 spinal cord segments at T11 vertebral level, L4-L5 at T12 vertebral level, and sacral segments at L1 vertebral level could be used for correlation (*7*, *36*). In our study, we were able to, for the first time reconstruct spinal cord segments from C2 to L5 using correlations between intervertebral foramen and rostral rootlet distance and vertebral bone length, along with segments lengths (Fig. 1A). With this approach, a potential implantation location of electrode array on spinal cord can be determined by mapping individual segments and related roots, using available imaging techniques, and later determine the optimal combination of the electrode leads for EES (Fig. 6). Previously, we used a similar approach on a swine model where segment lengths and vertebral bone lengths were found to have the strongest correlation at L2 level. This was translated into procedural electrode placement and further helped to optimize the electrode placement, demonstrating higher evoked motor responses while stimulating DREZ within segments compared to stimulating between the segments (*13*). Although variations in vertebral column and spinal cord anatomy between swine and human have to be considered, our present study demonstrates similar results and provides a fast and precise navigation approach, as only two parameters need to be correlated to reconstruct the entire spinal cord neuroanatomy. As bone structures grow more rapidly than the spinal cord during development after birth (*20*), which is more relevant for bipedal human that quadrupedal animals, adjustment based on the roots length (intervertebral foramen to rostral and caudal rootlet distances) helped to cover this variation.

## Conclusion

Understanding the spinal cord neuroanatomy and variations in basic anatomical measurements is critical for further development of SCS paradigms and for variety of surgical procedures. DREZ surgery has proven to be an effective intervention for relieving intractable pain of diverse etiologies (*19*) and SCS of the DREZ demonstrated capacity to alleviate chronic pain (*5*) and improved motor functions after SCI (*3*). Although, known fact that variation in spinal roots orientation has impact on outcome of EES (*8*), until now, there was no anatomy-based information available for human. Results of this study are important for performing surgical procedures on the spinal cord (*28*, *29*) and for implantation of epidural electrodes with later selection of optimal electrode contacts. Another finding of this study is that inter-root and inter-rootlet anastomoses could be considered for surgical and for neuromodulation procedures as they can be implicated in clinical sensory variations in nerve injury. Particular anatomic differences described in this study may help to further optimize spinal surgical procedures such as rhizotomy and drezotomy, improve the guidance when introducing the lesion-making device and for the location of spinal segments after laminectomy (*16*). These results may also facilitate a better understanding and management of postoperative complications (*14*-*17*).

## Materials and Methods

### Study Design

All study procedures were conducted with the approval of the Mayo Clinic Institutional Review Board (IRB) and in accordance with the National Institutes of Health (NIH) guidelines. Nine adult cadavers (5 males and 4 females) were used for this study with a mean age of 90.1 years (range 86-94), mean weight 63.1 kg (range 45.3-85.2), mean height 162.9 cm (range 150-192), and all of Caucasian ethnicity. The cadavers were fixed in a mixture of alcohol and formaldehyde before dissection.

### Spine Dissection and Anatomical Measurements

All cadavers were placed in prone position and intrinsic back muscles and ligaments of cervical, thoracic, and lumbar spine were carefully dissected. The vertebral column was harvested en bloc between C2-L5 in each cadaver. Once extracted, anatomical landmarks for vertebral bone were established (Table 1) and were measured manually using slide calipers. Then laminectomy was performed across all vertebrae to expose the spinal cord. The dura was incised to expose the spinal cord and main landmarks were determined and measured with slide calipers (Table 2). Next, high resolution pictures of each segment were taken with a surgical microscope (Leica M20, 4 x objective). Afterwards, the full bone dissection was performed and complete spinal cord was detached from the bone. Macroscopic pictures of entire spinal cord and at cervical, thoracic, and lumbar levels were taken followed by microscopic images of each spinal segment. The spinal cord was turned to its ventral side and dura was incised. Macroscopic pictures of ventral spinal cord were also taken similarly for cervical, thoracic, and lumbar levels, followed by microscopic image of each ventral segment. Dorsal and ventral microscopic images were analyzed using GeoGebra open source software to measure main spinal cord landmarks as well as rostral and caudal root angles. Number of dorsal and ventral rootlets were also counted using microscope.

**Table 2.** Anatomical measurements

| Dorsal and ventral roots | Spinal cord structures | Vertebral landmarks |
| --- | --- | --- |
| Rostral root angle | Transverse diameter | Mid-vertebral foramen length |
| Caudal root angle | DREZ length (rostral to caudal rootlet length) |  |
| Number of rootlets | Segment length at dorsal column and bone entry | Vertebral bone length |
| Width across columns | Inferior articular facet to caudal rootlet distance | Intervertebral foramen distance |
| Root diameter | Intervertebral foramen to rostral and caudal rootlet distance | Intervertebral foramen diameter |

### Schematic representation of dorsal segments mapping

A schematic representation of the spine and spinal cord for all measured segments (means of C2-L5) was done by plotting the following anatomic parameters to reconstruct a spine and spinal cord design to map specific segments: a) vertebral bone length, b) intervertebral foramen diameter, c) intervertebral foramen to rostral rootlet distance, d) intervertebral foramen to caudal rootlet distance, e) rostral root angle, f) caudal root angle, g) spinal cord transverse diameter, h) width across dorsal columns, i) DREZ length, and j) segment lengths at bone entry (Fig. 6C). Highest spinal cord correlation (intervertebral foramen distance to rostral rootlet distance to vertebral bone length) was used to stablish segmental relationship between vertebral bone landmarks and spinal cord (Fig. 6C, D).

### Data Analysis

Data Analysis: SigmaPlot (Systat Software, San Jose, CA, USA) was used to perform statistical analysis. The Shapiro-Wilk method was used to determine if the data were normally distributed. If so, an Equal Variance Test was performed using the Brown-Forsythe method. Significant differences were determined by one-way repeated-measures analysis of variance (ANOVA). If significance differences were detected pairwise multiple comparisons (Holm-Sidak) were performed to compare spinal segments. Data that was not normally distributed or variances that were not equal were analyzed using Kruskal-Wallis method based on ranks and pairwise multiple comparisons were done using Dunn’s method. Nine cadavers used for this study were acquired post-mortem following unrelated experimental studies in which the spinal column and cord remained intact.

### Correlation

Correlation analysis between spinal and spinal cord was done via Spearman correlation between the following spinal cord parameters: intervertebral foramen to rostral/caudal rootlet distance, mid-vertebral foramen length, vertebral bone length, and intervertebral foramen distance for all measured spinal cord segments C2-L5 (Fig. 6), in order to establish an anatomical segmental relationship in order to target dorsal spinal structures based on vertebral bone landmarks.

## Acknowledgments

The authors would like to thank body donors from the Mayo Clinic’s anatomical bequest program.

## Author contributions

A.M., R.I., and I.A.L., designed the study. A.M., R.I., T.L., O.J.J., B.K., P.S., and L.J.S., collected data. A.M., R.I., P.B., A.M.S., and P.J.G., analyzed the data. A.M., and R.I., created the figures. A.M., R.I., and I.A.L., wrote the manuscript text. N.L., and A.J.W., revised it. All authors reviewed the manuscript.

## Data and materials availability

All data associated with this study are present in the paper or the supplementary materials.

## References and Notes

1. C. A. Angeli, V. R. Edgerton, Y. P. Gerasimenko, S. J. Harkema. Altering spinal cord excitability enables voluntary movements after chronic complete paralysis in humans. Brain. 137, 1394–1409 (2014).

2. P. J. Grahn, I. A. Lavrov, D. G. Sayenko, M. G. Van Straaten, M. L. Gill, J. A. Strommen, J. S. Calvert, D. I. Drubach, L. A. Beck, M. B. Linde, A. R. Thoreson, C. Lopez, A. A. Mendez, P. N. Gad, Y. P. Gerasimenko, V. R. Edgerton, K. D. Zhao, K. H. Lee. Enabling Task-Specific Volitional Motor Functions via Spinal Cord Neuromodulation in a Human With Paraplegia. Mayo Clin Proc. 92, 544–554 (2017).

3. A. M. Mishra, A. Pal, D. Gupta, J. B Carmel. Paired motor cortex and cervical epidural electrical stimulation timed to converge in the spinal cord promotes lasting increases in motor responses. J. Physiol. 595, 6953–6968 (2017).

4. M. L. Gill, P. J. Grahn, J. S. Calvert, M. B. Linde, I. A. Lavrov, J. A. Strommen, L. A. Beck, D. G. Sayenko, M. G. Van Straaten, D. I. Drubach, D. D. Veith, A. R. Thoreson, C. Lopez, Y. P. Gerasimenko, V. R. Edgerton, K. H. Lee, K. D. Zhao. Neuromodulation of lumbosacral spinal networks enables independent stepping after complete paraplegia. Nat Med. 24, 1677–1682 (2018).

5. J. Holsheimer. Which Neuronal Elements are Activated Directly by Spinal Cord Stimulation. Neuromodulation. 5, 25–31 (2002).

6. B. Coburn, W. K. Sin. A theoretical study of epidural electrical stimulation of the spinal cord--Part I: Finite element analysis of stimulus fields. IEEE Trans Biomed Eng. 32, 971–7 (1985a).

7. F. Rattay, K. Minassian, M. R. Dimitrijevic. Epidural electrical stimulation of posterior structures of the human lumbosacral cord: 2. quantitative analysis by computer modeling. Spinal Cord. 38, 473–89 (2000).

8. B. Coburn. A theoretical study of epidural electrical stimulation of the spinal cord--Part II: Effects on long myelinated fibers. IEEE Trans Biomed Eng. 32, 978–86 (1985b).

9. J. Holsheimer, J. J. Struijk. How do geometric factors influence epidural spinal cord stimulation? A quantitative analysis by computer modeling. Stereotact Funct Neurosurg. 56, 234–49 (1991).

10. J. J. Struijk, J. Holsheimer, H. B. Boom. Excitation of dorsal root fibers in spinal cord stimulation: a theoretical study. IEEE Trans Biomed Eng. 40, 632–9 (1993).

11. J. Holsheimer. Computer modelling of spinal cord stimulation and its contribution to therapeutic efficacy. Spinal Cord. 36, 531–40 (1998).

12. J. Ladenbauer, K. Minassian, U. S. Hofstoetter, M. R. Dimitrijevic, F. Rattay. Stimulation of the human lumbar spinal cord with implanted and surface electrodes: a computer simulation study. IEEE Trans Neural Syst Rehabil Eng. 18, 637–45 (2010).

13. C. A. Cuellar, A. A. Mendez, R. Islam, J. S. Calvert, P. J. Grahn, B. Knudsen, T. Pham, K. H. Lee, I. A. Lavrov. The role of functional neuroanatomy of the lumbar spinal cord in effect of epidural stimulation. Front Neuroanat. 11, 82 (2017).

14. K. Shinomiya, A. Okawa, K. Nakao, K. Mochida, H. Haro, T. Sato, S. Heima. Morphology of C5 ventral nerve rootlets as part of dissociated motor loss of deltoid muscle. Spine. 19, 2501–4 (1994).

15. C. H. Jr. Alleyne, C. M. Cawley, D. L. Barrow, G. D. Bonner. Microsurgical anatomy of the dorsal cervical nerve roots and the cervical dorsal root ganglion/ventral root complexes. Surg Neurol. 50, 213–8 (1998).

16. J. P. Xiang, X. L. Liu, Y. B. Xu, J. Y. Wang, J. Hu. Microsurgical anatomy of dorsal root entry zone of brachial plexus. Microsurgery. 28, 17–20 (2008).

17. A. A. Kumar, S. Seshayyan, V. Tamilalagan, M. Sindou. Surgical anatomy of dorsal root entry zone of cervical spinal nerves: Cadaveric study. Int J Anat Res. 2, 296–04 (2014).

18. A. Karatas, S. Caglar, A. Savas, A. Elhan, A. Erdogan. Microsurgical anatomy of the dorsal cervical rootlets and dorsal root entry zones. Acta Neurochir. 147, 195–9 (2005).

19. M. Bozkurt, S. Canbay, G. F. Neves, E. Aktüre, E. Fidan, M. S. Salamat, M. K. Başkaya. Microsurgical anatomy of the dorsal thoracic rootlets and dorsal root entry zones. Acta Neurochir. 154, 1235–9 (2012).

20. J. H. Kim, C. W. Lee, K. S. Chun, W. H. Shin, H. G. Bae, J. C. Chang. Morphometric Relationship between the Cervicothoracic Cord Segments and Vertebral Bodies. J Korean Neurosurg Soc. 52, 384–90 (2012).

21. M. W. Zhou, W. T. Wang, H. S. Huang, G. Y. Zhu, Y. P. Chen, C. M. Zhou. Microsurgical anatomy of lumbosacral nerve rootlets for highly selective rhizotomy in chronic spinal cord injury. Anat Rec. 293, 2123–8 (2010).

22. T. Mamatha, S. Madhyastha, M. D. Prameela, B. V. Murlimanju, V. Saralaya. Morphology of nerve rootlets and spinal segments in the lumbosacral region: An anatomical study. Research Journal of Pharmaceutical, Biological and Chemical Sciences. 7, 1848–1854 (2016).

23. Y. Liu, X. Zhou, J. Ma, Y. Ge, X. Cao. The diameters and number of nerve fibers in spinal nerve roots. J Spinal Cord Med. 38, 532–537 (2015).

24. J. S. Calvert, P. J. Grahn, J. A. Strommen, I. A. Lavrov, L. A. Beck, M. L. Gill, M. B. Linde, D. A. Brown, M. G. Van Straaten, D. D. Veith, C. Lopez, D. G. Sayenko, Y. P. Gerasimenko, V. R. Edgerton, K. D. Zhao, K. H. Lee. Electrophysiological Guidance of Epidural Electrode Array Implantation over the Human Lumbosacral Spinal Cord to Enable Motor Function after Chronic Paralysis. J. Neurotrauma. 35, 1–10 (2018).

25. M. Capogrosso, N. Wenger, S. Raspopovic, P. Musienko, J. Beauparlant, L. Bassi Luciani, G. Courtine, S. Micera. A computational model for epidural electrical stimulation of spinal sensorimotor circuits. J Neurosci. 33, 19326–40 (2013).

26. G. Kayalioglu. The Spinal Cord. Chapter 4 – The Spinal Nerves (2009).

27. W. Pallie. The Intersegmental Anastomoses of Posterior Spinal Rootlets and Their Significance. J Neurosur. 16, 188–196 (1959).

28. R. S. Tubbs, D. EL-Zammar, M. Loukas, A. Cömert, A. A. Cohen-Gadol. Intradural cervical root adjacent interconnections in the normal, prefixed, and postfixed brachial plexus. J Neurosurg Spine. 11, 413–416 (2009).

29. B. Solmaz, N. Tatarlı, D. Ceylan, E. Keleş, S. Çavdar. Intradural communication between dorsal rootlets of spinal nerves: their clinical significance. Acta Neurochir. 157, 1069–76 (2015).

30. J. M. Marzo, E. H. Simmons, F. Kallen. Intradural connections between adjacent cervical spinal roots. Spine. 12, 964–8 (1987).

31. N. Tanaka, Y. Fujimoto, H. S. An, Y. Ikuta, M. Yasuda. The Anatomic Relation Among the Nerve Roots, Intervertebral Foramina, and Intervertebral Discs of the Cervical Spine. Spine. 25, 286–291 (2000).

32. J. Moriishi, K. Otani, K. Tanaka, S. Inoue. The intersegmental anastomoses between spinal nerve roots. Anat Rec. 224, 110–116 (1989).

33. M. M. Pinter, F. Gerstenbrand, M. R. Dimitrijevic. Epidural electric stimulation of posterior structures of the human lumbosacral cord: 3. Control of spasticity. Spinal Cord. 38, 524–31 (2000).

34. M. R. Dimitrijevic, Y. Gerasimenko, M. M. Pinter. Evidence for a spinal central pattern generator in humans. Ann N Y Acad Sci. 860, 360–376 (1998).

35. J. Lang, U. Geisel. Über den lumbosakralen teil des durasackes und die topographie seines inhalts. Morphol Med. 3, 27–46 (1983).

36. E. J. Wall, M. S. Cohen, J. J. Abitbol, S. R. Garfin. Organization of intrathecal nerve roots at the level of the conus medullaris. J Bone Joint Surg Am. 72, 1495–9 (1990).

